# Expression of cell-adhesion molecules in *E. coli*: a high-throughput method to identify paracellular modulators

**DOI:** 10.1101/2021.04.08.439041

**Authors:** Jay Rollins, Tyler Worthington, Emily Hooke, Joseph Hobson, Jacob Wengler, Sandra Hope, Dario Mizrachi

## Abstract

Cell-adhesion molecules (CAM) are membrane proteins responsible for cell-cell interactions or cell-extracellular matrix interactions. Among these proteins, claudins (CLDN), occludin (OCLN), and junctional adhesion molecules (JAM) are components of the tight junction (TJ), the single proteic structure tasked with safeguarding the paracellular space. The TJ is responsible for controlled permeability of blood-tissue barriers, regulating the passage of molecule passage by size and charge. Currently there is no translational solution to manipulate the TJ with the exception of Focused Ultra-sound (FUS) and Micro bubbling (MB) techniques, still in clinical trials. Here we describe the expression of TJ proteins in the outer membrane of *E. coli* and report its consequences. When expression is induced, the unicellular behavior of *E. coli* is replaced with multicellular aggregations that can be quantified using Flow Cytometry (FC). The adhesion properties of the aggregates are representative of the individual membrane proteins expressed. This method, called iCLASP (inspection of cell-adhesion molecules aggregation through FC protocols), allows the high-throughput interrogation of small-molecules influence on paracellular permeability, enabling for the first time the discovery of its modulators for therapeutic strategies.

## Introduction

Cell adhesion molecules (CAMs) are proteins located on the cell surface involved in binding with other cells or with the extracellular matrix (ECM) in a process called cell adhesion^1^. CAMs are complex membrane proteins. In the classical sense, CAMs are grouped into four major families: selectins, immunoglobulin superfamily (IgSF), integrins, and cadherins. Recently, novel integral membrane proteins, claudins (CLDN) and occludin (OCLN), have been identified as major cell adhesion molecules working at the tight junction (TJ)^2^. Claudins comprise a multigene family, and each member of approximately 23 kDa bears 4-α-helix transmembrane domains^3^. OCLN is a unique protein of the TJ with structural homology to CLDN’s 4-α-helix transmembrane domains^4^. Junctional adhesion molecules (JAM), another membrane component of the TJ, is a member of the IgSF^5^. CAMs involvement in developmental and physiological processes^6, 7^, or pathophysiological events such as tumorigenesis and metastasis^8, 9^, enhances their relevance in translational solutions^10^.

Epithelial and endothelial cell-cell contacts are needed for homeostasis as intercellular junctional complexes are key for their maintenance. Among them, the adherens junctions (AJs) provide essential adhesive and mechanical properties. The TJs occupy the most apical region of the cell and create an almost impenetrable barrier that forms without interruption in the entire perimeter of the cell. AJs and TJs play essential roles in vascular permeability, but only the TJ controls the paracellular permeability^11^.

As an example of the relevance of TJs we will examine briefly the blood-brain barrier (BBB). The BBB is a cellular barrier that maintains the homeostasis of the neural microenvironment^12^. The TJs between brain capillary endothelial cells greatly limit molecules to traffic across the paracellular route, with the exception of small molecules (<500 Da) and gaseous molecules^13^. Additionally, brain transcytosis occurs at low rates as compared with peripheral tissues, restraining the vesicle-mediated transcellular transport of macromolecules^14^. The paracellular and transcellular barrier properties of BBB also set up challenges for drug delivery to the central nervous system (CNS). Current solutions to overcome the BBB in the clinical setting are extremely limited. The BBB prevents approximately 98% of small molecule drugs from entering the brain^15, 16^. Focused ultrasound (FUS) combined with intravenous microbubbles (MBs) is a promising technique, being both temporary and reversible, that increases BBB permeability ^17^. FUS induced BBB permeability is reestablished within 24 h^18^. Side effects caused by FUS and MBs can be detected by MRI. FUS-induced edema has been reported^19^. MRI images have detected extravasation of red blood cells and are used to evaluate vascular damages following FUS treatment^20^. A review of FUS and MBs was published recently^21^. Data demonstrates that with these treatments the degree of BBB permeability is increased while the risk of tissue damage also rises, in some cases with months-long side effects^22, 23^.

TJ’s dysfunction in the BBB is responsible for increased permeability^24^. Thus, controlling the TJ in the BBB can result in advantageous manipulation of the paracellular permeability. The development and delivery of small molecule drugs is relatively straightforward. Drug discovery of small molecules from target selection through to clinical evaluation is a very complex are challenging areas of drug discovery. The main obstacle is the initial hit-finding. For example, the combination of so little effort in developing solutions to the BBB permeability leads directly to the present situation in neurotherapeutics with few effective treatments for the majority of brain-related disorders. This situation can be reversed by an accelerated effort, knowledge base in the fundamental transport properties of the BBB, and the molecular and cellular biology of the brain capillary endothelium. Currently, dynamics of the BBB and cytotoxicity testing and drug permeation are carried out *in vitro* using Trans epithelial electrical resistance (TEER)^25^. Some challenges encountered in TEER are lengthy set-up, up to three weeks prior to obtaining measurements, and the availability of suitable cell lines that represent the desired system of study. Finally, identified cellular systems may fail to produce adequate and measurable TEER values^25, 26^.

Prodrug methods used to improve drug penetration via the transcellular pathway have been successfully developed, and some prodrugs have been used to treat patients. The use of transporters to improve absorption of some drugs (e.g., antiviral agents) has also been successful in treating patients. Other methods, including (a) blocking the efflux pumps to improve transcellular delivery and (b) modulation of cell-cell adhesion in the intercellular junctions to improve paracellular delivery across biological barriers are still in the investigational stage.

Using a synthetic biology approach, we engineered *E. coli* to recombinantly express TJ membrane proteins. Our method, named iCLASP (<u>i</u>nspection of <u>c</u>ell-<u>a</u>dhesion molecule<u>s</u> aggregation through FC <u>p</u>rotocols) is a high throughput solution to identify paracellular modifiers to potentiate drug delivery of hydrophilic molecules.

## Results and Discussion

### Expression system design

The initial goal was to recombinantly express adhesion molecules in the outer membrane of *E. coli*. We decided to create a fusion protein between OmpW (accession number P0A915) and a human CLDN. The N- and C-terminus of outer membrane proteins are located in the periplasm thus a fusion between OmpW and CLDN will result in an exposure of the adhesive domains (extracellular loops) to the exterior of the cell (Figure 1.A). A challenge to this design was the recombinant expression of JAM-A. JAM proteins have an N-terminus that contains the adhesive immunoglobulin domains, followed by the transmembrane domain. To overcome this bottleneck, we converted OmpW to a circularly permutated protein (cpOmpW) after the design offered for OmpX, another *E. coli* protein^27^. The cpOmpX^27^ protein and our cpOmpW create new N- and C-terminus that are now located outside of the cell (Figure 1.B). This strategy enabled the fusion of cpOmpW and JAM-A. The linker between the fused proteins contains a TEV protease cleavage site. Treatment of cells for 2-hours at room temperature releases cpOmpW from JAM-A and enables the interpretation of the adhesive properties of JAM-A.

**Figure 1.**
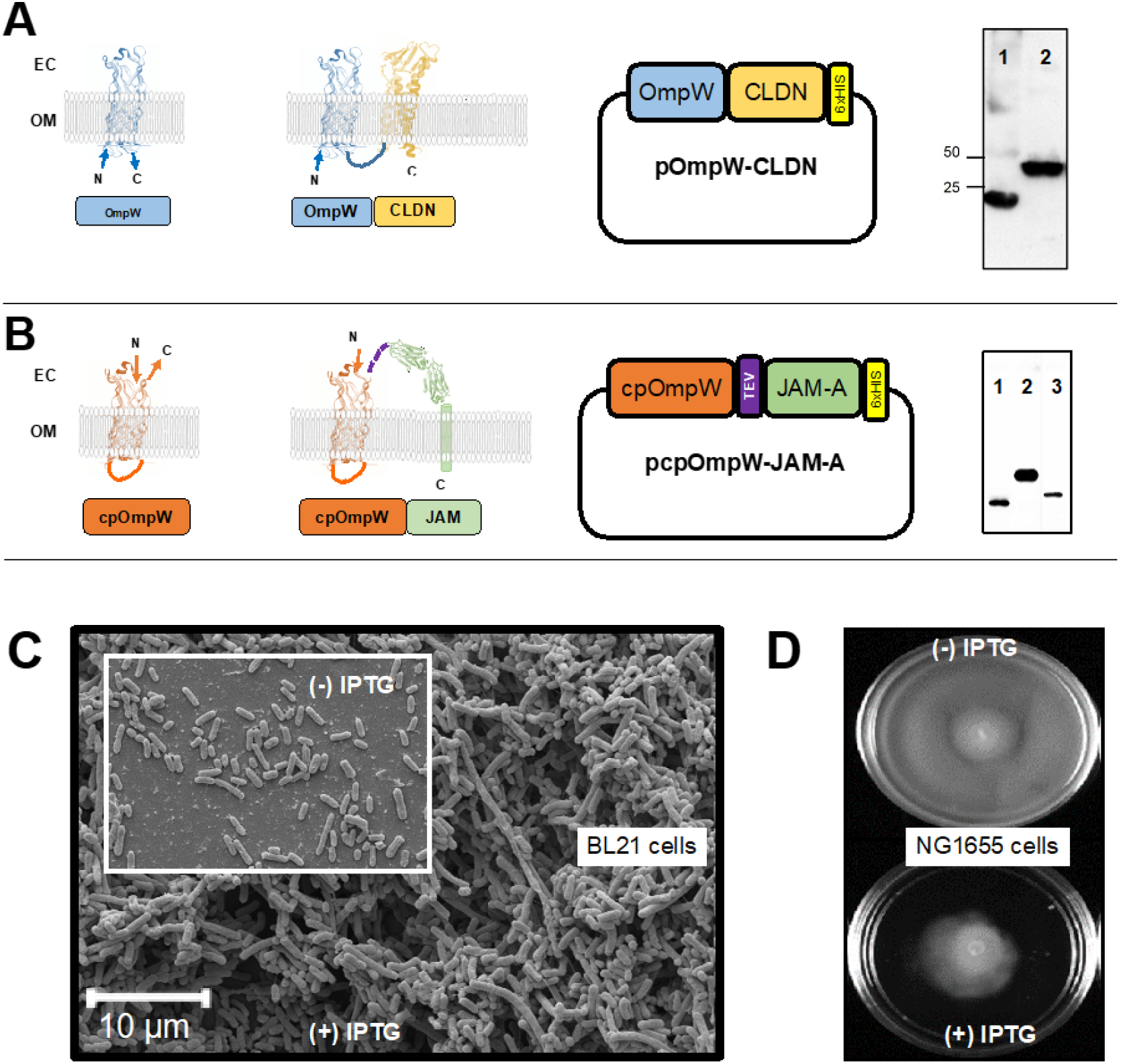
Strategy for CAM expression in *E. coli.* **A)** OmpW is a native outer membrane protein of *E. coli* that populates the outer membrane, its N- and C-terminus are located in the extracellular space. These features make OmpW a suitable fusion partner for 4-α-helix CAMs like CLDN, OCLN, and others. Plasmids are synthesized with a C-terminal His-tag for Western blot detection with anti-His antibody (Abcam, product number ab1187, Cambridge, MA, USA): 1-OmpW (24 kDa),2-OmpW-CLDN (47 kDa). **B)** Circularly permutated OmpW (Supplementary file) is a suitable fusion partner for CAM proteins, which have an extracellular N-terminus. This strategy was employed to recombinantly express JAM-A. A linker region containing TEV protease sequence enables for separation of cpOmpW and JAM-A post expression. The Plasmid contains a His-tag C-terminal to JAM-A. A Western blot with anti-His antibody is shown: 1-cpOmpW, 2-cpOmpW-JAM-A, 3- cpOmpW-JAM-A after TEV protease cleavage (only JAM-A is observed). **C)** Consequences of overexpressing CLDN in *E. coli*. BL21 DE3cells are captured in Electron Microscopy (http://www.ihcworld.com/_protocols/em/em_negative2.htm) displaying their natural behavior (unicellular) in a Negative Staining experiment (insert). In the background photo, BL21 DE3 after 16 hours of protein expression of OmpW-CLDN, in the images the large aggregates are evident. **D)** NG1655, a swimming variant of *E. coli*, growing on plates of LB+0.25% agar. Top image contains NG1655 cells after 24 hours of growth, from a single drop cells at OD_600_=1, placed at the center of the plate. In contrast, and under identical conditions, when protein expression (OmpW-CLDN) is induced (1 mM IPTG in the LB+ agar plate), cells growth is reduced, after 24 hours. The plate demonstrates that aggregation of the cells, induced by expression of OmpW-CLDN, prevents them from displaying their typical swimming behavior, and cannot advance beyond a small radius from the center.

### Flow Cytometry of bacterial cells expressing TJ membrane proteins

BL21 DE3 cells are transformed with the corresponding plasmids (CLDNs, OCLN, or JAM-A). Cells grow to an OD600 of 1, induced with 1 mM IPTG and allowed to continue to grow for 18 hours at Room Temperature. Cells are prepared for 96-well plates setup. Samples were prepared in suspension (50 μL cells per well and 150 μL of PBS) and run through a Beckman Coulter Cytoflex flow cytometer (Beckman Coulter, Indianapolis, IN, USA). Readings were collected using the side scatter (SSC) data from the 405 nm (violet) laser for excitation and a 405/10 bandpass filter for emission detection. The violet laser SSC has a greater sensitivity than the forward scatter (FSC) or SSC detection of the 488 nm (blue) laser for detecting alterations in cell shape as reported by others^28^ and according to our assessment of flow data results in this study. SSC area versus height readings were plotted for data analysis. Cytoflex-generated FCS flow data files were analyzed using FlowJo 10 software (BD Biosciences, Ashland, OR, USA). The violet SSC-area by SSC-height data was gated for data analysis set to exclude upper and lower extremes that would interfere with the calculation of the slope of the line (Figure 2). Structural cell changes are detected as the area readings move away from the height readings where area increases at a lower rate than height and the slope of the line decreases (Figure 3). Further manipulation of the data using R Studio software, generously prepared by Stephen Picollo, Biology Department, Brigham Young University, resulted in Experimental Slopes calculated for each sample (Materials and Methods) and used to plot the results.

**Figure 2.**
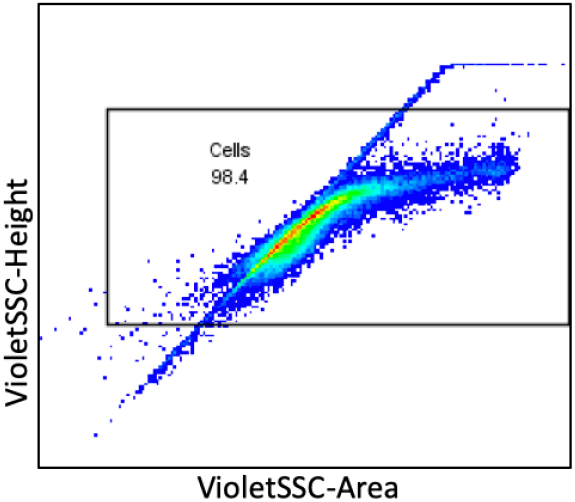
Gating of area and height readings. Data was gated prior to determining the slope of the line to eliminate the upper and lower ends of the data. At least 98% of the readings were included in the gate for analysis.

**Figure 3.**
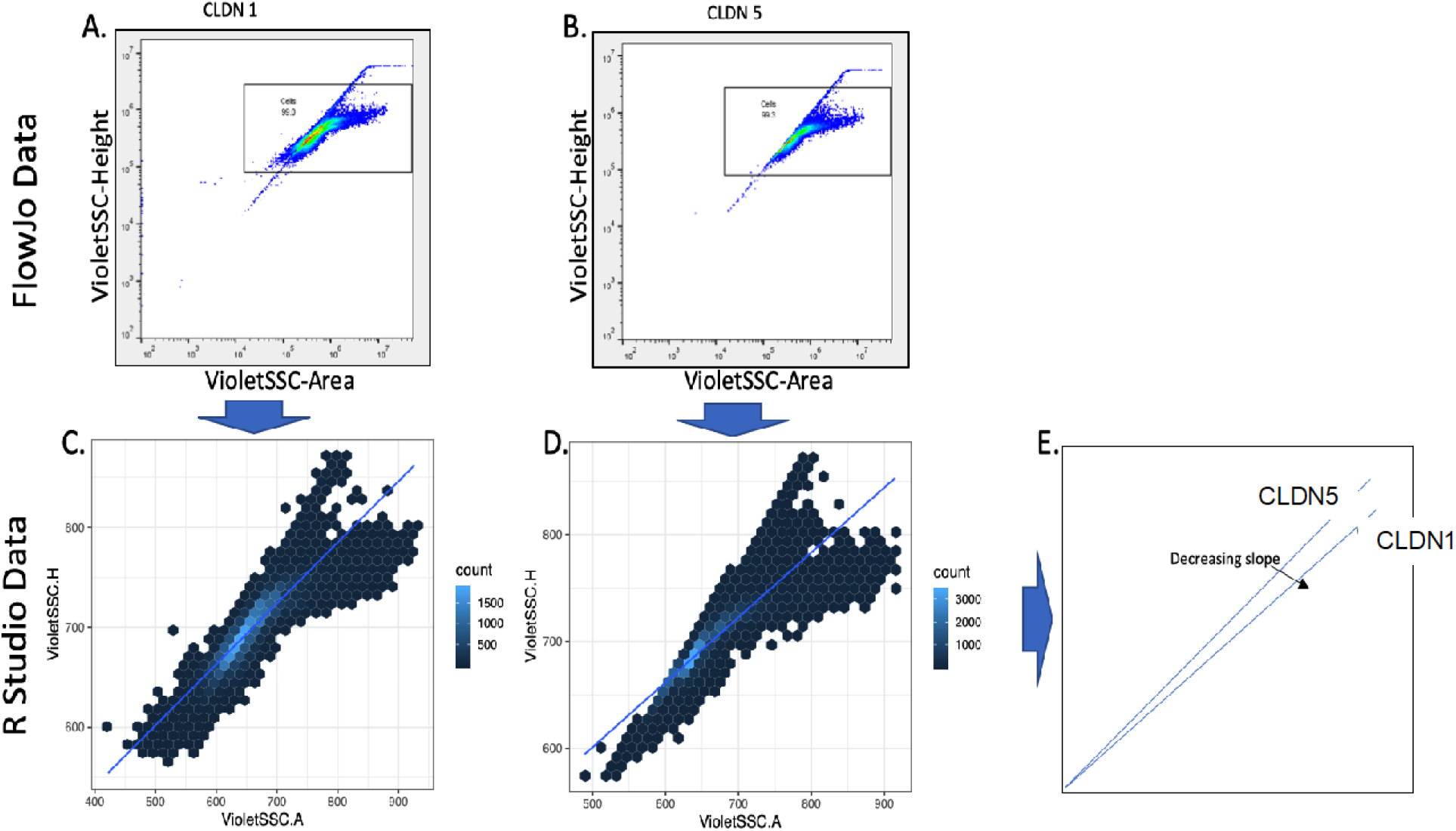
Data analysis using FlowJo and R Studio software. Flow cytometry data was gated in FlowJo software (panels A and B) and R Studio converted the data into Hexbin graphs (panels C and D) and determine the slope of the line (panel E).

Endowed with this tool we proceeded to study the adhesive properties of TJ proteins. CLDNs are a family of 27 proteins in mammals^3^. We used OmpW as a fusion partner to express CLDN1 through 10, OCLN, and tricellulin (TRCL). The latter is the first integral membrane protein found to concentrate at the vertically oriented TJs of tricellular contacts.^29^. We included in the analysis stargazing (STRGZ), an AMPA receptor^30^ believed to have some structural and functional homology to CLDN. Figure 4 correlates the adhesive properties of the above-mentioned proteins by plotting the Experimental Slopes calculated as described. The control for these experiments are BL21 DE3 cells with plasmid pET28a, and BL21 DE3 cells recombinantly expressing OmpW alone in a pET28a backbone.

**Figure 4.**
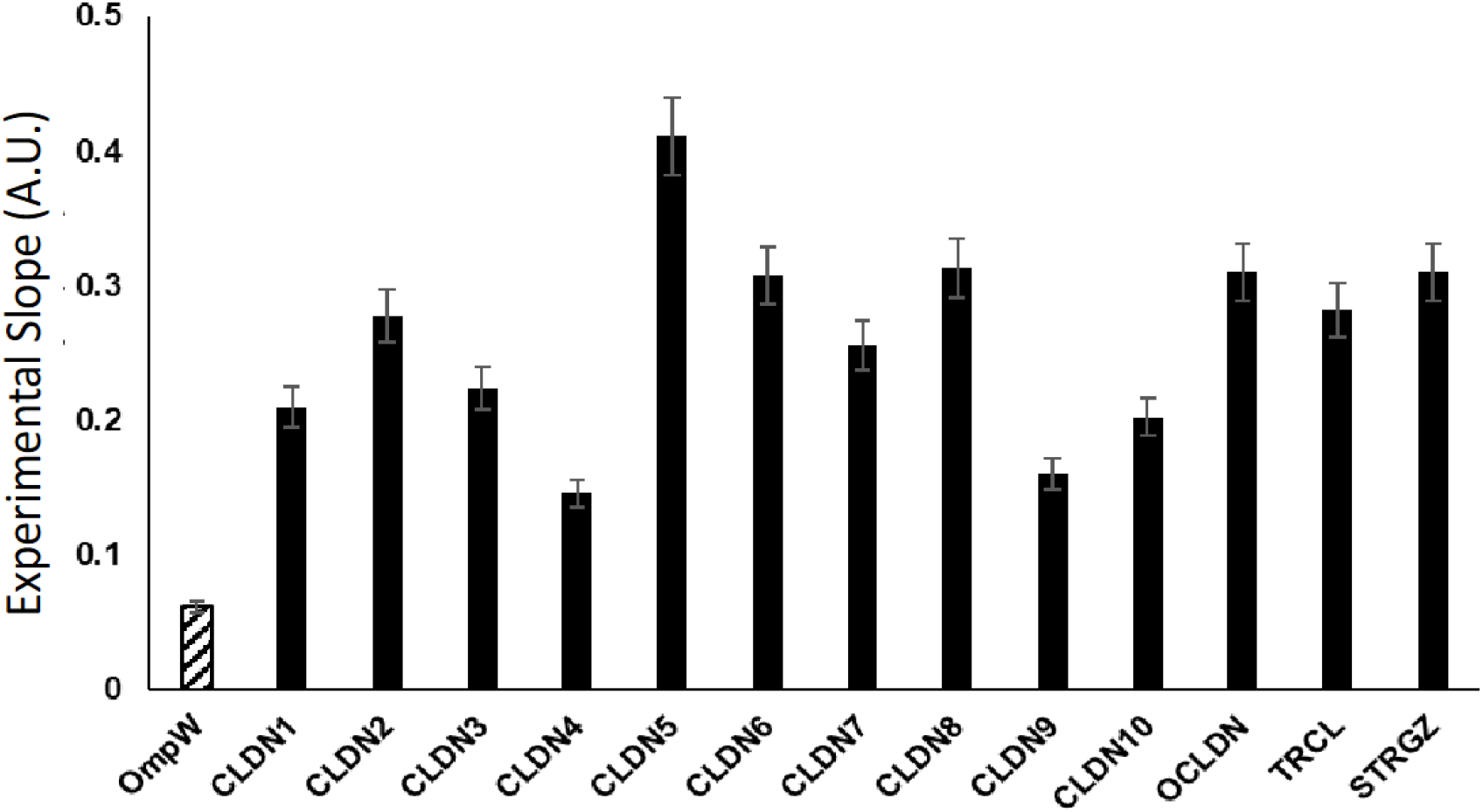
Experimental slopes of 4-α-helical CAMs. Protein expression of 4-α-helical CAMs is presented here. Flow Cytometry analysis, described in this article, enables the determination of Experimental Slopes that represent adhesive properties of CAMs. Data presented here was statistically analyzed. All points are expressed as the average value of 12-replicates in 4 different experiments (n=4) ± SE. All points are statistically significant and significantly different from each other. Asterisks are omitted for display purposes. Additionally, all values presented are the result of subtracting the value of BL21 DE3 cells with pET28a (0.92±0.039).

To determine the extent at which iCLASP can be used to determine changes in TJ-induced paracellular permeability we resourced to study a dose response of molecules known to non-specifically alter TEER. Sodium Caprate is a known detergent that disrupts CLDN-CLDN interactions resulting in increased permeability^31^. On the other hand, ethanol has been described as an agent that increases CLDN-CLDN interactions, resulting in a decreased permeability^32^.

Figure 5 demonstrates dose-dependent changes to permeability caused by Caprate or Ethanol. Our data suggests iCLASP is capable of identifying changes to CLDN-CLDN and also OCLN-OCLN interactions and can be used for discovery of small molecules that affect paracellular permeability controlled by TJs.

**Figure 5.**
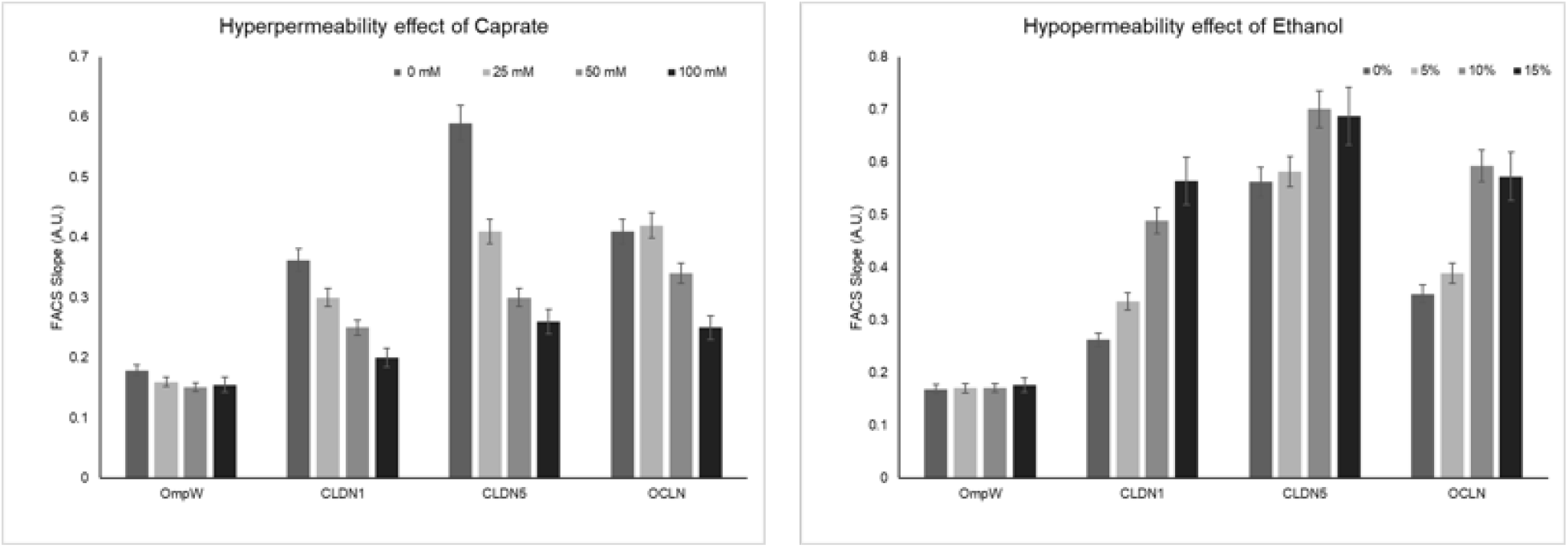
Experimental slopes of 4-α-helical CAMs in the presence of paracellular permeability influencers. Protein expression of 4-α-helical CAMs or OmpW alone, is presented here. Flow Cytometry analysis is plotted here to demonstrate the dose-dependent effect of Caprate (left panel), an agent know to disrupt paracellular permeability. The right panel displays the dose-dependent effect of ethanol, a known agent to increase tightness of the paracellular space. The effects of Caprate and ethanol are observed in CLDN1, CLDN5 and OCLN but not on OmpW alone. Data presented here was statistically analyzed. All points are expressed as the average value of 12-replicates in 4 different experiments (n=4) ± SE. All points are statistically significant and significantly different from each other, at all concentrations of the influencer. Asterisks are omitted for display purposes. Additionally, all values presented are the result of subtracting the value of BL21 DE3 cells with pET28a (0.92±0.039).

### Ion permeable CLDNs and the Hofmeister effect

CLDNs are key regulators of barrier properties of the TJ. CLDNs are better recognized for their primary responsibility to tighten the paracellular pathway. A few CLDNs, and therefore the TJs they regulate, serve as paracellular channels for water and ions^33^. Among CLDNs with these properties we have selected cation-selective human CLDN2^34^ and anion-selective human CLDN10^35^.

The Hofmeister series characterizes the ions as to their ability to “salt-in” or “salt-out” proteins. Experimental results show that ion cooperativity may play an important role in affecting water properties^36, 37^. In Figure 6, both CLDN2 and CLDN10 are examined under the effects of slats that have been well characterized in the Hofmeister Series^38–40^. We employed the same salts for CLDN1, CLDN2 and CLDN10 (see Materials and Methods). CLDN1 did not display any trends under conditions of any of the salts employed. Chloride salts (Al^+3^, Mg^+2^, Na^+^, and Rb^+^) only show a trend with CLDN2 (Figure 5). Sodium salts in the series representing different negative groups (HPO_4_^−^, Cl^−^, NO_3_^−^, and ClO_4_^−^) only show a trend with CLDN10 and no other CLDNs tested (Figure 5).

**Figure 6.**
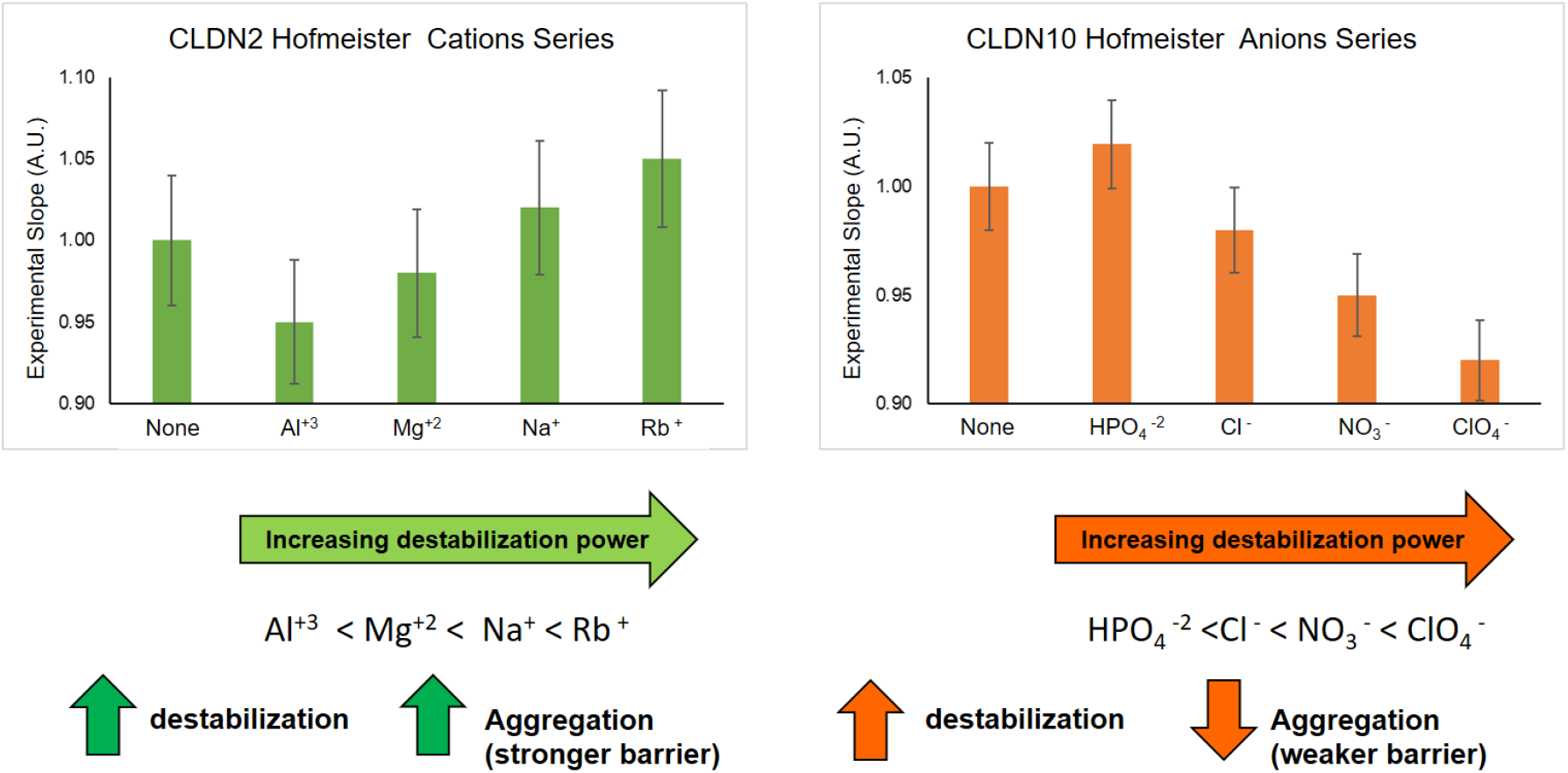
The Hofmeister series and ion permeable CLDNs. Ion permeable human CLDN2 (cations) and human CLDN10 (anions) were exposed to the influence of different salts. For CLDN2 Chloride salts were used. For CLDN10 Sodium salts were used. The left panel indicates the Hofmeister disruptive effects of cations. Similarly, the right panel displays the Hofmeister ranking of disruptive effects of the negative ions. Flow Cytometry analysis enables the determination of Experimental Slopes that represent adhesive properties of CAMs in the presence of 100 mM salts. Data presented here was statistically analyzed. All points are expressed as the average value of 12-replicates in 4 different experiments (n=4) ± SE. All points are statistically significant and significantly different from each other. Asterisks are omitted for display purposes.

For CLDN2, the Hofmeister series for cations has a trend of increasing cell-cell adhesion (Figure 5). Al^+3^ having a more relaxing effect on CLDN2-CLDN2 interactions, while Rb^+^ fosters increased CLDN2-CLDN2 interaction. The Hofmeister series ranks the salts as stabilizing or the degree of destabilization. These rankings are denoted in Figure 5. In the case of CLDN2, losing stability drives further CLDN2-CLDN2 interactions. This effect could be detrimental to the cation permeability function, as it will tighten the TJ but reduce ability to sort cations. CLDN10, on the other hand, has a trend that seems to indicate that at a lower stability the permeability of the TJ will increase. Taken together, when salts decrease stability of the microenvironment, CLDN2 loses function but tightens the TJ (hypopermeability), while CLDN10 loses function and the TJ’s organization that can lead to hyperpermeability.

### iCLASP for the TJ’s JAM-A

As described above (Figure 1.B) the iCLASP method was slightly modified to accommodate JAM-A, a very important adhesion molecule where the N-terminus is located in the extracellular domain, opposite to CLDN, OCLN, and other 4-α-helical TJ membrane proteins where the N-terminus is intracellular. By circularly permutation of OmpW (cpOmpW) we were able to fuse JAM-A such that its adhesive domains are exposed to the extracellular space. A cleavable linker (TEV protease) was placed between cpOmpW and JAM-A to observe the adhesion properties of JAM-A as a fused protein or as a free molecule on the surface of *E. coli*. BL21 DE3 cells hosting the plasmids empty, cpOmpW or cpOmpW-JAM-A were grown over night after induction as described above. Each group is further divided in two: no treatment or TEV protease treatment (5 U/mL) for 2-hours at room temperature. Further, cells are diluted 1:4 with PBS and 200 μL are dispensed per well in a 96-well plate for Flow Cytometry. Four different experiments were carried out in 12-replicates each. The data displayed in Figure 7 describes the ability of iCLASP to successfully identify an increase of cell-cell interactions when BL21 DE3 recombinantly expressed cpOmpW-JAM-A. No significant difference was observed after TEV treatment. This could be a unique case of JAM-A and its quaternary structure^41^ to foster cell-cell adhesion. In Figure 1.B we show a Western blot that indicates that after 2-hour of TEV protease treatment the cleavage is successful. Other CAMs with a similar secondary structure (immunoglobulin domains) to that of JAM-A can also be studied using iCLASP. We expect that in other cases the treatment could be more significant and will need to be tailored for the corresponding CAM.

**Figure 7.**
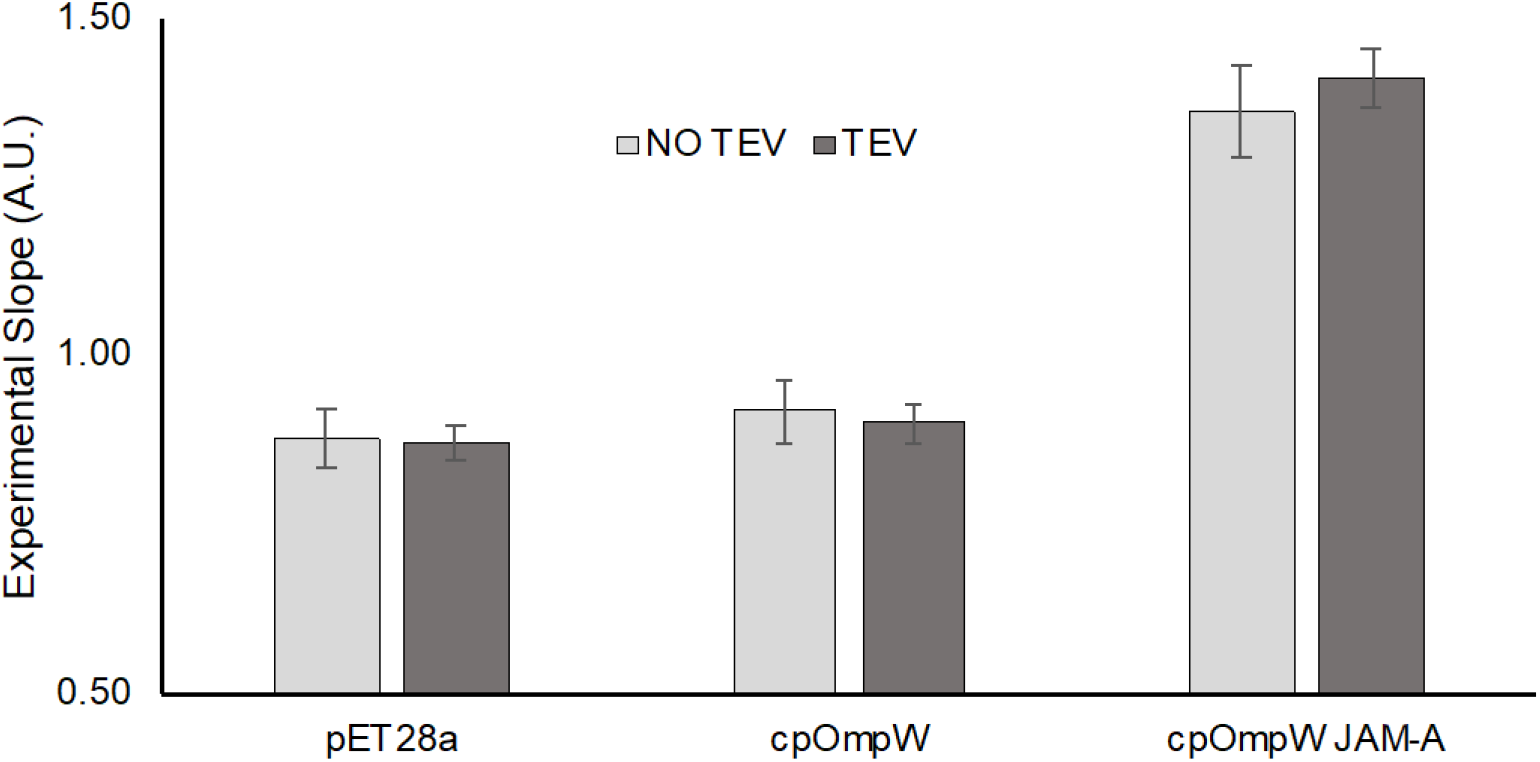
Flow Cytometry analysis of JAM-A fused to cpOmpW. Flow Cytometry analysis is resented here for the recombinantly expression of cpOmpW-JAM-A protein. Experimental slopes of empty plasmid (pET28a), cpOmpW protein alone and the fusion with JAM-A are plotted in the graph. The higher Experimental Slope indicates the presence of BL21 DE3 cells aggregates, which represent the adhesive properties of JAM-A. Data presented here was statistically analyzed. All points are expressed as the average value of 12-replicates in 4 different experiments (n=4) ± SE. All points are statistically significant and significantly different from each other. Asterisks are omitted for display purposes.

### iCLASP model

Finally, we propose a graphical model to illustrate the power of iCLASP (Figure 8).

**Figure 8.**
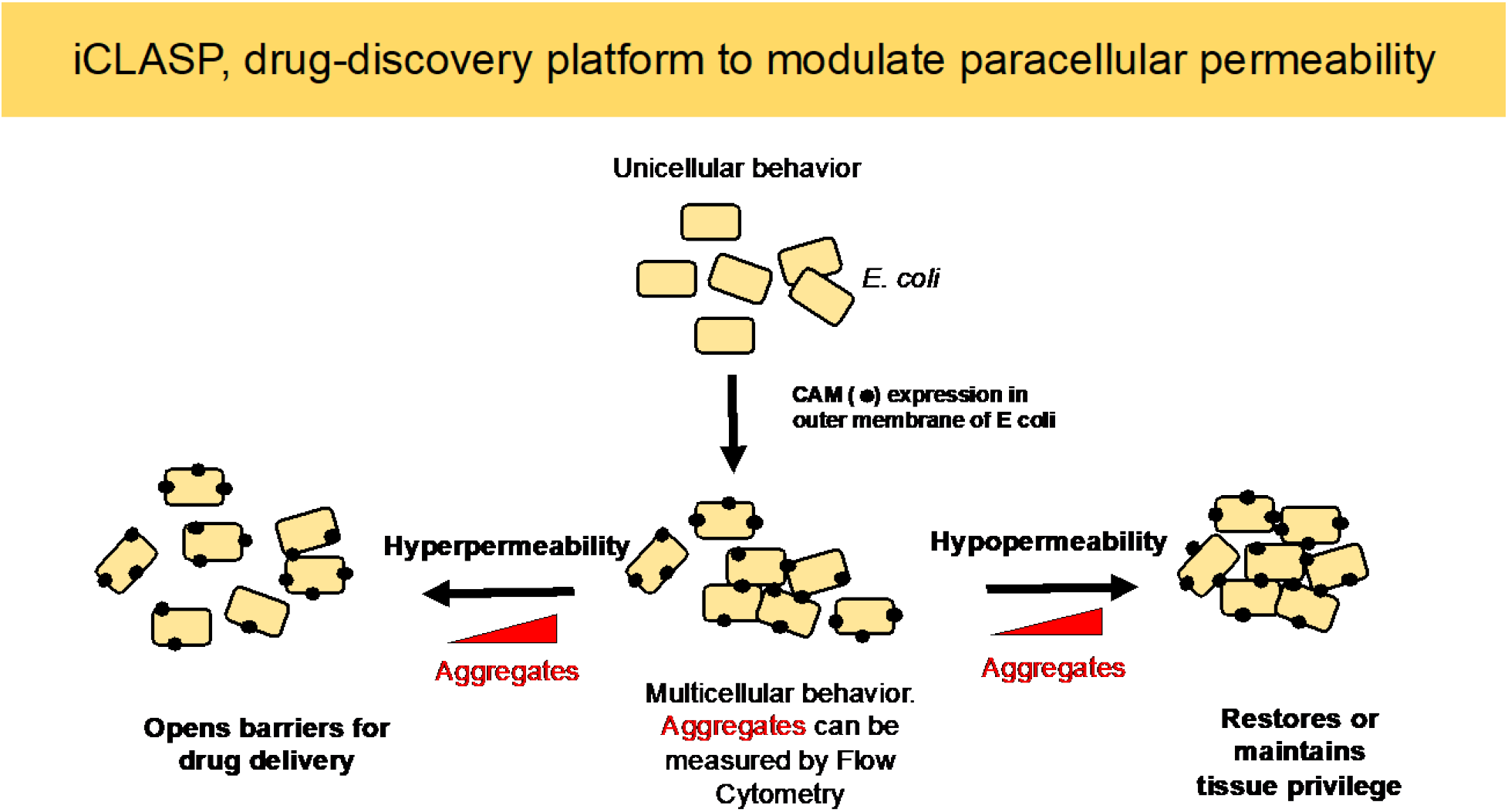
Conceptual Model. The iCLASP experimental model is presented here.

Bacterial cells are transformed with plasmids hosting CAMs, regardless of their secondary structure or orientation in the mammalian plasma membrane. CAMs induce multicellular behavior in *E. coli* (aggregates or clumps of cells) that can be measured by Flow Cytometry protocols. A typical experiment will contain cells expressing CAMs untreated, as internal control accounting for growth and protein expression variability; and treated. The power displayed by iCLASP is two-fold, the screen of a library of small compounds may identify, in the same experiment, compounds that decrease or increase the size of aggregates. The first will represent cases in which the compound induces hyperpermeability of the paracellular space, while the second represents a hypopermeability outcome. Hyperpermeability may be desirable when trying to overcome, as an example, the BBB for drug delivery. TJ dysfunction may contribute to epithelial permeation disorder and multiple intestinal diseases like inflammatory bowel diseases (IBD)^42^. In such cases, hypopermeability may be required to foster cell-cell interactions to prevent progression of the disease.

## Conclusion

Currently, there are no therapeutic solution to control the permeability of the paracellular space. CAMs like the TJ’s integral membrane proteins are responsible for paracellular permeability. In this article we present evidence for a high throughput *E. coli*-based method that recombinantly expresses CAMs. This expression results in bacterial cell aggregates that can be inspected using Flow Cytometry. The method, named iCLASP, can examine small molecule libraries and identify candidates that increase or decrease permeability. Compared to classical methods, iCLASP is faster, only 4 days to set up (from transformation to experiment), and Flow Cytometry is performed at speeds of one 96-well plate every 45 minutes. CAMs are responsible for a number of other cell-cell, cell-extracellular matrix, cell-pathogen interactions. The iCLASP method offers a model to examine CAMs behavior in a cellular environment that is often challenging in mammalian cells.

## Materials and Methods

### Reagents and Genes

For Flow Cytometry, flat bottom 96-well cell culture plates were employed (GeneClone, https://geneseesci.com/). All genes employed in this study were synthesized by TWIST biosciences (https://twistdna.com, San Francisco, CA, USA) and cloned in pET28a. All salts used for experimental performance of Figure 6 were obtained from Sigma Aldrich (https://www.sigmaaldrich.com/, St. Louis, MO, USA). TEV protease was obtained from New England Biolabs (Ipswich, MA, USA).

### Transformation, Cell growth, and Protein Expression (LB, fresh transformations, controls)

Plasmids (pET28a, pET28a-OmpW-CLDN, pET28a-cpOmpW-JAM-A) are transformed in BL21 DE3 cells, plates are prepared LB, 2% agar, and 100 ug/mL of kanamycin. A single colony is grown over night in 5 mL of LB and kanamycin. A 1:1000 dilution of the overnight culture is started in the morning. Cells are grown at 30 °C until OD_600_ is between 0.9-1.0. IPTG (1 mM) is used to induce protein expression, cells are placed in shaker at 21 °C (room temperature) and allowed to continue growth for 18 hours. Samples can be analyzed by Western blot and anti-HIS antibody.

### Flow Cytometry data collection and analysis

Samples were prepared in suspension for analysis and run through a Beckman Coulter Cytoflex flow cytometer (Beckman Coulter, Indianapolis, IN, USA). Readings were collected using the side scatter (SSC) data from the 405 nm (violet) laser for excitation and a 405/10 bandpass filter for emission detection. The violet laser SSC has a greater sensitivity than the forward scatter (FSC) or SSC detection of the 488 nm (blue) laser for detecting alterations in cell shape as reported by others^28^ and according to our assessment of flow data results in this study. SSC area versus height readings were plotted for data analysis. Cytoflex-generated FCS flow data files were analyzed using FlowJo 10 software (BD Biosciences, Ashland, OR, USA). The violet SSC-area by SSC-height data was gated for data analysis sets to exclude upper and lower extremes that would interfere with the calculation of the slope of the line. Structural cell changes are detected as the area readings move away from the height readings where area increases at a lower rate than height and the slope of the line decreases. Below is the code used for R Studio software analysis, generously prepared by Stephen Picollo, Biology Department, Brigham Young University. Code lines with # indicate the functions of the code. The code uses a working file from FlowJo, generates hexbin plots to visualize analyzed data, and generates a spreadsheet of the calculated slopes as a .csv file.

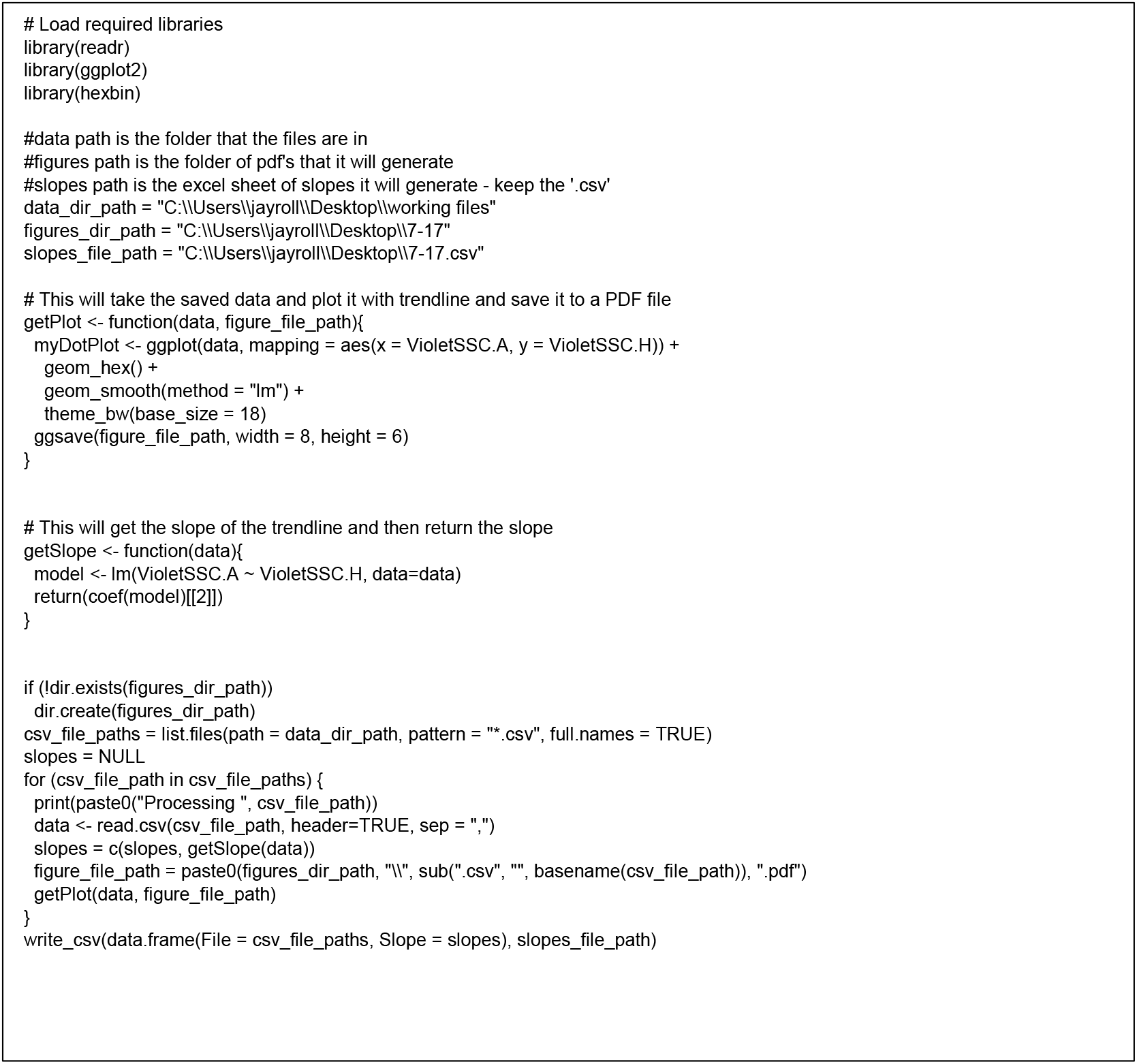

### Statistical Analysis

Flow Cytometry data (Experimental Slope) were analyzed using SAS software (SAS Institute Inc., Cary, NC, United States) and the Mixed Procedure method to generate p-values, standard deviation, and standard error and to determine statistical significance (for Figure 5). For all experiments α = 0.05. Data was collected for each sample in four different experiments (n=4). Each condition was measured in 12-replicates. Thus, each data points corresponds to the average of 12-replicates and n=4. Statistical differences were identified for all samples in each graph. The final analysis concluded that all treatments are statistically significant (p<0.0001) and significantly different from each other and the control (asterisks omitted for display purposes).

## Acknowledgements

Stephen Piccolo prepared R Studio software code to determine the slope of the line from data in FlowJo.

## Supplementary material

Amino acid sequences and DNA sequences of genes used in this study

### 1. pET28a- OmpW (His tag). Cloned between NcoI and XhoI

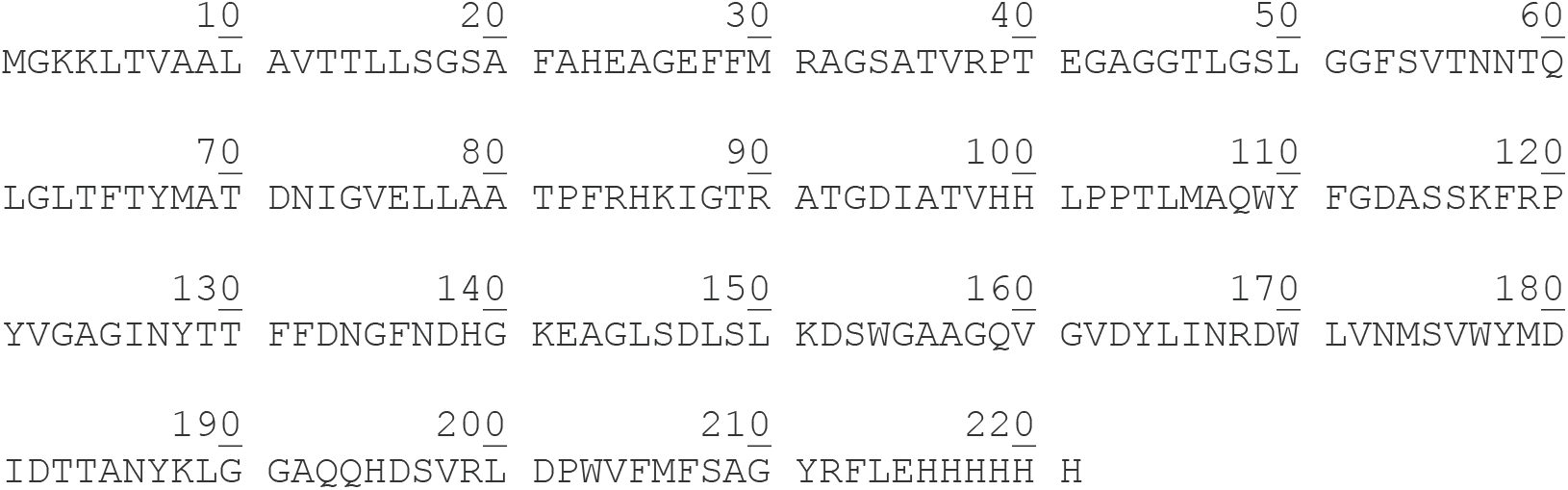

### 2. pET28a- OmpW-hCLDN1 (His tag). Entire fusion cloned between NcoI and XhoI, an NdeI site was placed in amino acids 219 and 220 for easy substitution with other targets (NdeI toXhoI)

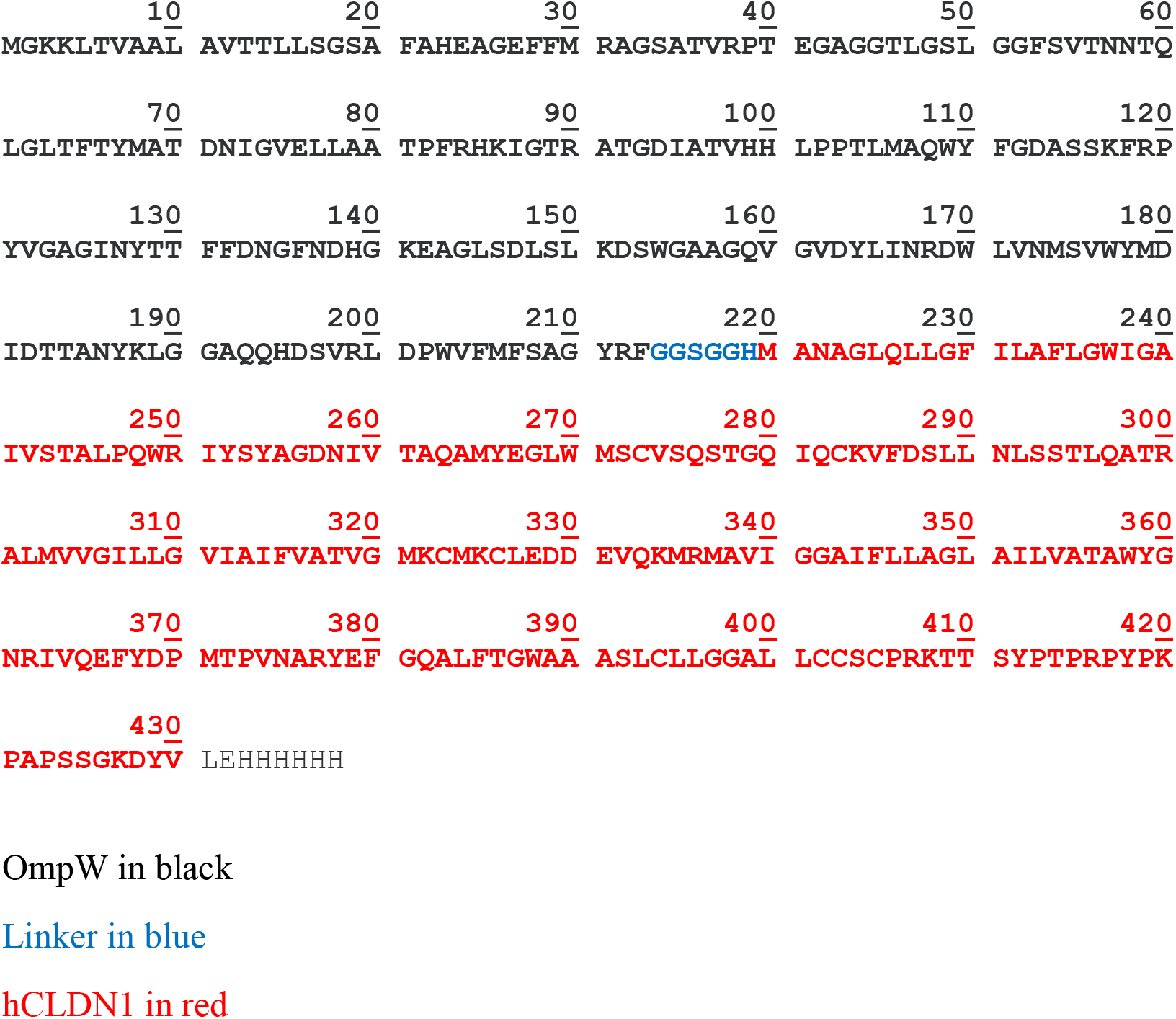

### 3. pET28a- cpOmpW-TEV (His tag)

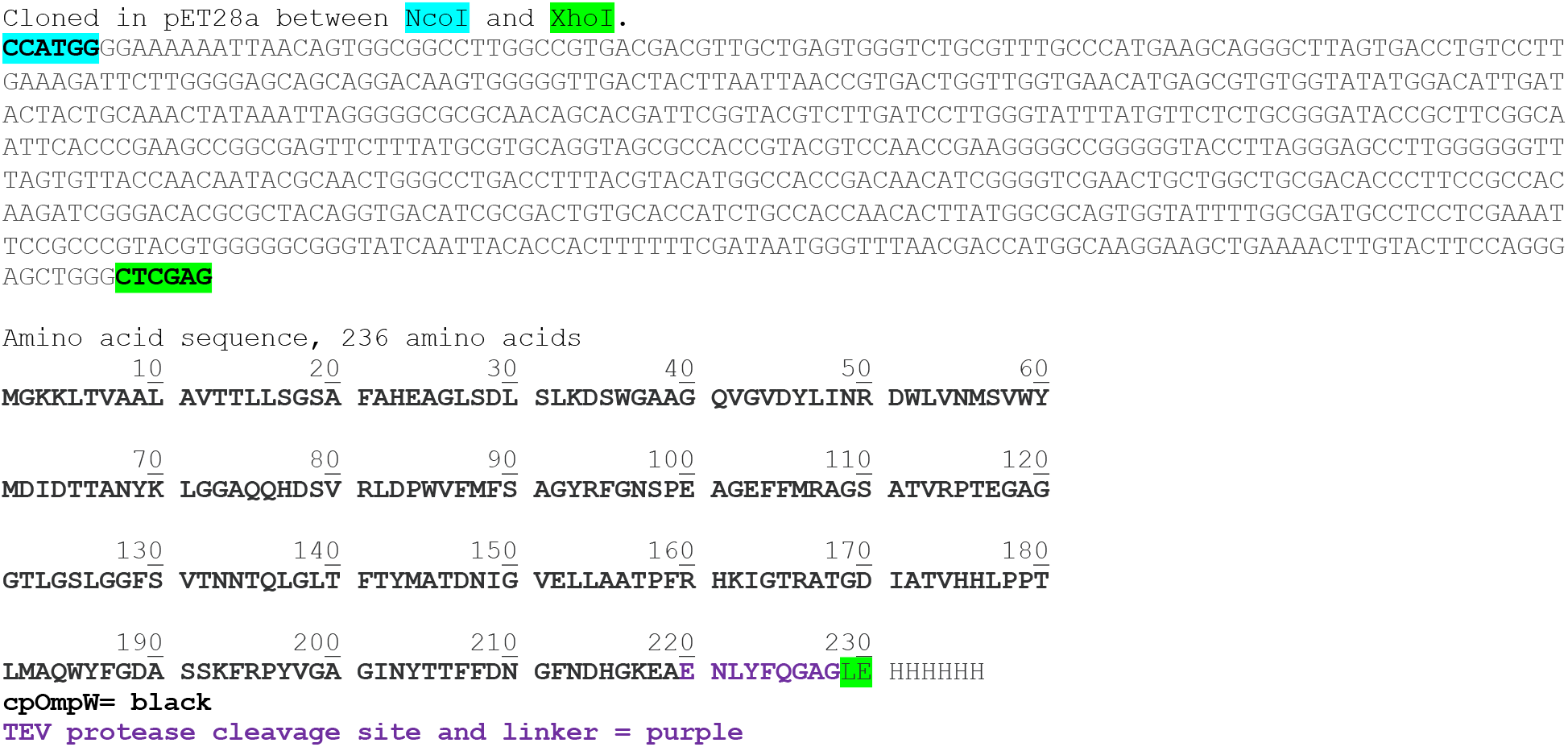

### 4. pET28a- cpOmpW-TEV-hJAM-A (His tag)

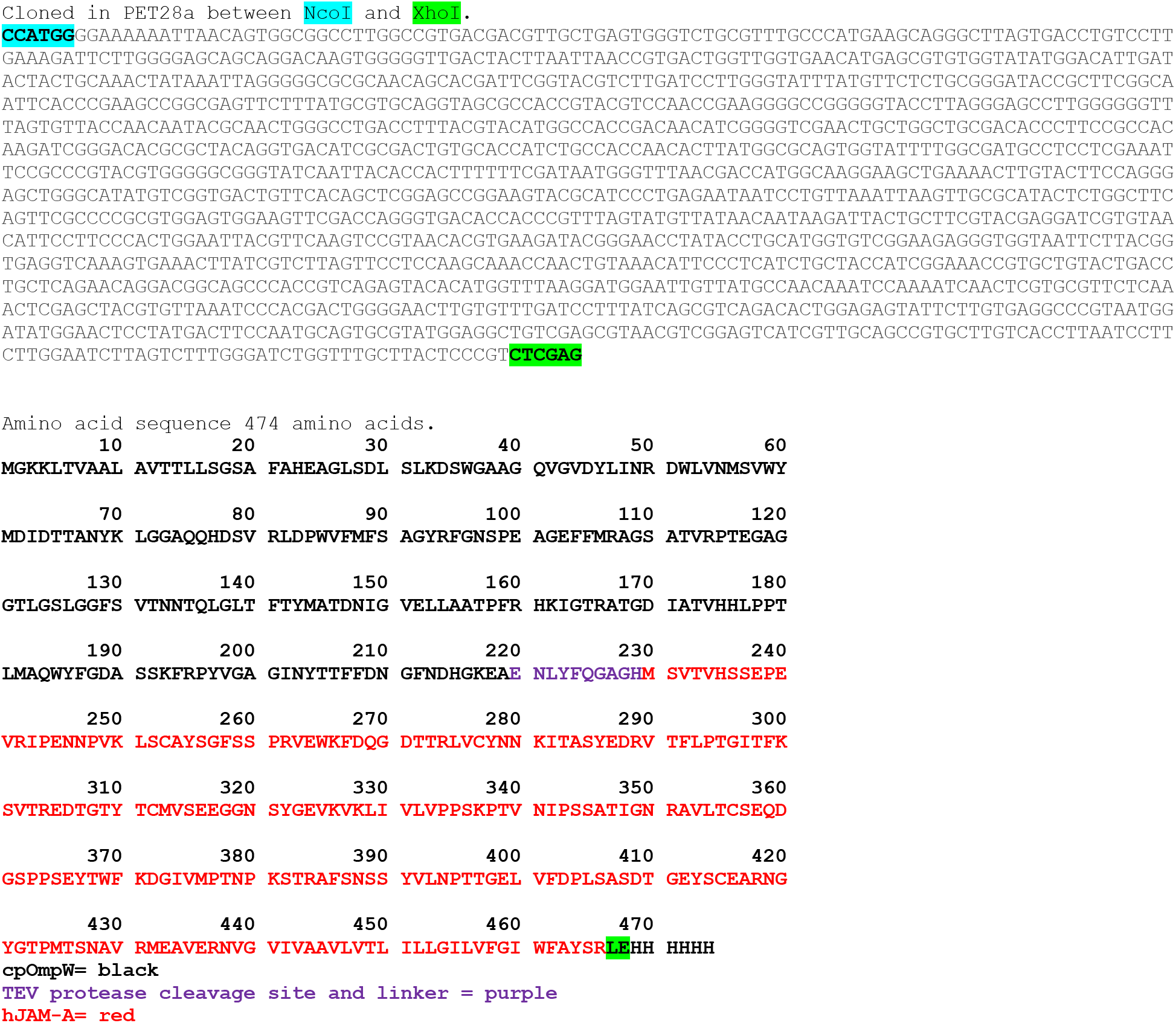

Proteins in this study and their Accession numbers. Proteins were used full length unless specified.

| Protein | Accession number | URL |
| --- | --- | --- |
| Human CLDN1 | O95832 | https://www.uniprot.org/uniprot/O95832 |
| Human CLDN2 | P57739 | https://www.uniprot.org/uniprot/P57739 |
| Human CLDN3 | O15551 | https://www.uniprot.org/uniprot/O15551 |
| Human CLDN4 | O14493 | https://www.uniprot.org/uniprot/O14493 |
| Human CLDN5 | O00501 | https://www.uniprot.org/uniprot/O00501 |
| Human CLDN6 | P56747 | https://www.uniprot.org/uniprot/P56747 |
| Human CLDN7 | O95471 | https://www.uniprot.org/uniprot/O95471 |
| Human CLDN8 | P56748 | https://www.uniprot.org/uniprot/P56748 |
| Human CLDN9 | O95484 | https://www.uniprot.org/uniprot/O95484 |
| Human CLDN10 | P78369 | https://www.uniprot.org/uniprot/P78369 |
| Human Tricellulin<br>(TRCL) | Q8N4S9<br>(Met186-His368) | https://www.uniprot.org/uniprot/Q8N4S9 |
| Human Stargazin<br>(STRGZN) | P62955<br>(Met1-Arg203) | https://www.uniprot.org/uniprot/P62955 |
| Human JAM-A | Q9Y624 | https://www.uniprot.org/uniprot/Q9Y624 |

